# TRPV4 Promotes Histamine Receptor Signaling in Lymphatic Endothelial Cells

**DOI:** 10.1101/2025.11.25.690563

**Authors:** Jeffri S Retamal, Scott Peng, Thanh-Nhan Nguyen, Esther Bandala-Sanchez, Emily M Eriksson, Roisin E McCague, Narges S Mahdavian, Simona E Carbone, Enyuan Cao, Arisbel B Gondin, Nicholas A Veldhuis, Daniel P Poole

**Author notes:** Co-corresponding authorship. Equal contribution (Jeffri S Retamal, Scott Peng). **Funding sources** National Health & Medical Research Council Australia APP2021163 (NAV, ABG) and APP2021675 (DPP, SEC); Australian Research Council (ARC) Future Fellowship FT220100617 (NAV); ARC DECRA Fellowship DE200100825 (SEC), Australia Research Council Centre of Excellence in Convergent Bio-Nano Science and Technology.

## Abstract

**Background:** The control of lymphatic permeability and flow is essential for homeostatic regulation of tissue fluid balance and immune responses. Histamine has been identified as an important signaling mediator involved in the regulation of lymphatic function. Histamine is released from activated perilymphatic mast cells and may also be produced by lymphatic endothelial cells (LECs) in response to flow-induced shear stress. The non-selective cation channel Transient Receptor Potential Vanilloid 4 (TRPV4) is an important mediator of signaling by GPCRs, including histamine receptors. TRPV4 is activated in response to shear stress and is functionally expressed by LECs. We hypothesized that histamine receptors and TRPV4 interact in LECs, leading to activation of distinct downstream signaling pathways. This study examined the mechanistic link between TRPV4 activity and histaminergic signaling in LECs.

**Principle Results:** Histaminergic Ca^2+^ signaling was examined in primary human LECs. Responses to histamine were mainly driven by the H1R histamine receptor, with some contribution by the H4R subtype, as determined using selective antagonists. H4R signaling in response to 4-methylhistamine was effectively prevented by either removal of extracellular Ca^2+^ or block of TRPV4 activity, consistent with TRPV4-dependence. Conversely, activation of H4R resulted in marked sensitization of subsequent responses to the selective TRPV4 agonist GSK1016790A. This interaction was mediated through a PLA2-dependent mechanism. TRPV4 activity was required for histamine receptor-evoked translocation of the Ca^2+^-dependent transcription factor NFATc1 and for cytoskeletal remodeling. By contrast, the release of cytokines in response to activation of either histamine receptors or TRPV4 were largely independent processes.

**Conclusions:** This study identifies TRPV4 as an important mediator of histaminergic signaling in LECs. The findings provide further support for the involvement of TRPV4 in defining the nature and magnitude of endothelial signaling downstream of GPCRs.

**Highlights:**

- Histamine receptors are functionally expressed by primary human LECs
- TRPV4 is an important driver of H4R-evoked Ca^2+^ signaling in LECs
- Histamine receptor activation sensitizes TRPV4 signaling in LECs
- TRPV4 promotes histamine-evoked NFATc1 translocation to the nucleus of LECs
- Histamine and TRPV4 evoked cytokine release from LECs involve distinct mechanisms

**Graphical abstract:** Graphical abstract.
Histamine exerts its effects on lymphatic endothelial cells through interaction with the H1R and H4R receptor subtypes. H1R activation promotes Ca^2+^ release from intracellular stores. Activation of the H4R leads to elevated intracellular Ca^2+^ through crosstalk with the non-selective cation channel TRPV4. Interaction between H4R and TRPV4 is mediated through a PLA2-dependent mechanism. TRPV4 enhances histamine-evoked NFATc1 activation and translocation and cytoskeletal remodeling. In contrast, histamine receptor- and TRPV4-mediated cytokine release appear to involve mechanistically independent processes.

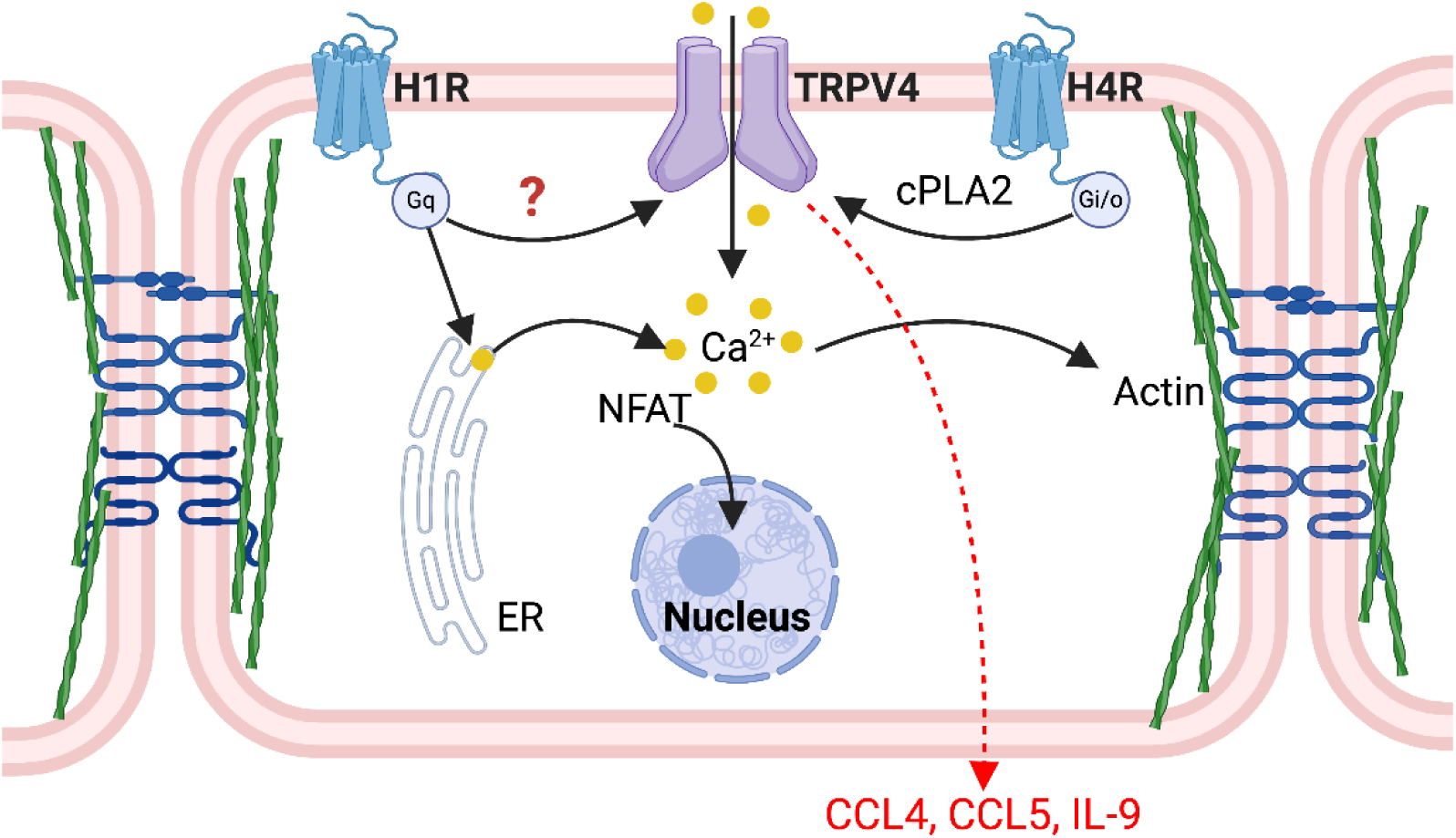

## 1. INTRODUCTION

### 1.1 Lymphatic system

The lymphatic vasculature performs critical functions including regulating fluid balance, lipid uptake, removal of waste, and immune surveillance. Disruption to lymphatic drainage due to developmental defects, damage, or disease, can lead to pathological changes including edema and inflammation [1]. The lymphatic system has established roles in a diverse range of diseases and conditions including inflammatory bowel disease [2], tumor metastasis [1], and cardiac disease [3].

The lymphatic system runs in parallel to the blood vasculature and is organized into initial lymphatic capillaries, pre-collecting lymphatic and collecting lymphatic vessels. Lymphatic capillaries located within tissues are structurally composed of a single layer of lymphatic endothelial cells (LECs) with loose ‘button-like’ junctions, leading to a comparatively high degree of permeability to cells and macromolecules relative to blood capillaries.

### 1.2 TRPV4 regulates lymphatic vessel function

Mechanosensitive ion channels including Piezo1 and Orai1 have an integral role in the development of the lymphatic vasculature [4, 5]. The importance of mechanosensory signaling for lymphatic development is exemplified by Piezo1 loss of function mutations, which impact valve formation and are associated with familial lymphedema [5–7]. LECs also express the non-selective cation channel Transient Receptor Potential Vanilloid 4 (TRPV4), which is a mechanotransducer involved in cellular responses to shear stress. Although TRPV4 influences lymphatic pumping and flow [8–10], the flow rate and associated shear stress in lymphatic vessels is significantly lower than that in blood vasculature ([11]: 0.9 mm/s; 0.4-0.6 dyn/cm^2^) and is below the threshold required for TRPV4 activation [12]. TRPV4 is also activated by other mechanisms including functional coupling to G protein-coupled receptors (GPCRs; [13]) and to Piezo1 [14]. GPCRs can activate or enhance TRPV4 signaling through several pathways including production of endogenous TRPV4 activating lipids and channel phosphorylation [15, 16]. The importance of GPCR-TRPV4 interactions for physiological and pathophysiological processes has been demonstrated for a broad range of GPCRs and systems [13]. This includes functional coupling of TRPV4 to signaling by 5-HT and protease-activated receptors in endothelial cells of blood vessels, leading to enhanced vascular leak [15, 17, 18]. Equivalent interactions are likely to be of functional importance for LECs and lymphatic vessels. However, comparatively limited information is available regarding the expression and function of GPCRs in LECs.

### 1.3 Histamine receptors functionally interact with TRPV4

Histamine is the primary endogenous ligand for specific histamine GPCRs, of which there are four subtypes (H1R-H4R), each with distinct coupling mechanisms [19, 20]. Prior evidence indicates that histamine promotes relaxation of lymphatic vessels through activation of the H1R and H2R subtypes [21]. Histamine is released from degranulated perilymphatic mast cells and is associated with reduced endothelial barrier function [22] and altered immune cell trafficking to lymphatic vessels [23]. Histamine may also be endogenously produced and released by LECs in response to shear stress and can influence lymphatic vessel function through an autocrine mechanism [24].

The potential for functional interaction between histamine receptors and TRPV4 has been demonstrated for H1, H2, and H4 receptors in cell types including sensory neurons [25, 26] and keratinocytes [27]. In this study we examine how TRPV4 influences the magnitude and duration of histamine receptor signaling in LECs. We identify the importance of TRPV4 as a primary driver of Ca^2+^ signaling by the H4 histamine receptor subtype (H4R). Furthermore, we demonstrate that activation of H4R can lead to sensitization of TRPV4 signaling. These findings place TRPV4 as a central mediator of histaminergic signaling in LECs, with implications for lymphatic function in both health and disease.

## 2. MATERIALS AND METHODS

### 2.1 Reagents

Reagents were obtained from the following suppliers: histamine, diphenhydramine, ranitidine, S38093-2, GSK1016790A and DAPI from Sigma-Aldrich (Castle Hill, NSW, Australia); 4-methylhistamine hydrochloride, arachidonyl trifluoromethyl ketone (AACOCF3), HC067047, ionomycin, phorbol 12,13-dibutyrate (PDBu), and YM-26734 from Cayman Chemical (Michigan, USA); Fura2-AM and JNJ7777120 from Abcam (Cambridge, UK).

### 2.2 Cell culture

Adult human dermal microvascular lymphatic endothelial cells (HMVEC-dLy) (Lonza #CC-2810) were cultured in endothelial cell growth medium-2 (Lonza #CC-3202) consisting of EBM-2 basal media and supplements provided in a bullet kit (containing: FBS (5%), hFGF-B (0.4%), VEGF (0.1%), R3-IGF-1 (0.1%), hEGF (0.1%), ascorbic acid (0.1%), hydrocortisone (0.04%), and GA-1000 (antibiotic mixture of 30µg/ml gentamicin + 15ng/ml amphotericin B; Lonza #CC-4083; 0.1%)). Cells were used for a maximum of 8 passages. Cells were grown in a humidified incubator at 37℃ and 5% CO_2_.

### 2.3 Calcium signaling assay

LECs were seeded onto non-coated 96-well plates (15,000 cells/well) and grown to ≥80% confluence. Cells were loaded with Fura2-AM ester (1.5µM) in calcium assay buffer (150mM NaCl, 2.6mM KCl, 1.2mM MgCl_2_, 2.2mM CaCl_2_, 10mM glucose, and 10mM HEPES in water) with probenecid (2mM) and pluronic acid (0.5µM) (pH adjusted to 7.4) for 30 min at 37℃. Loading buffer was removed and cells were washed with the same calcium assay buffer without Fura-2 and pluronic acid for 30 min. The Ca^2+^ chelator EGTA (2 mM), antagonists and inhibitors of signaling pathways were added at this point. Fluorescence was measured at 340/380 nm excitation and 510 nm emission wavelengths using a FlexStation 3 plate reader (Molecular Devices LLC, San Jose, CA, USA). Agonists were added after 20 s of baseline reading (1:5 ratio to calcium assay buffer). Measurements were taken every 4 s for a total of 150 s at 37℃. Maximal effects were determined based on peak Ca^2+^ responses by LECs to stimulation with ionomycin (1 µM). Data were processed using SoftMax Pro software (Molecular Devices).

### 2.4 Immunofluorescence and microscopy

LECs were seeded onto collagen-coated glass coverslips and cultured to ≥80% confluence. After treatments, the cells were washed, then fixed with 4% PFA in PBS for 15 min. Cells were blocked and permeabilized (PBS containing 0.1% saponin, 0.1% sodium azide, and 5% horse serum; 1 h, RT). Cells were incubated with a rabbit anti-VE-cadherin antibody (Abcam ab33168; RRID:AB_870662, 1:400) or mouse monoclonal antibody targeted to NFATc1-2 (clone 7A6, BioLegend 649602; RRID: AB_10679126, 1:500) in the same blocking buffer (overnight, 4°C). The next day, cells were washed (3x PBS), then incubated with a donkey anti-rabbit or anti-mouse Alexa Fluor 647 secondary antibody (ThermoFisher, 1:500) and Alexa Fluor 488-conjugated phalloidin (Abcam, ab176753, 1:1,000) in PBS (2 h, RT). Cell nuclei were labeled with DAPI (1:2,000 in PBS, 15 min, RT). Coverslips were mounted onto slides with ProLong Diamond antifade mountant (ThermoFisher). Cells were imaged using a Leica TCS-SP8 confocal system with an HC PLAN APO 1.4 NA 63x glycerol objective. Images were captured at 16-bit-depth and 1024×1024 pixel resolution. Images were presented as maximal Z-projections of 8-12 stacks (30 µm/stack).

### 2.5 Image analysis

Nuclear NFATc1-2 was analyzed using the Fiji distribution of ImageJ [28]. For each treatment, NFATc1-2 positive pixels were determined from 50 cells per group from 4 technical replicates. The nuclear-specific NFATc1-2 labeling was identified based on DAPI-labeling and expressed relative to the total NFATc1-2 positive pixels in the cell.

### 2.6 Bio-Plex™ Cytokine Assay

Cytokine secretion by LECs was quantified using Luminex xMAP technology (Bio-Plex™ 200, Bio-Rad). LECs were seeded into 96-well plates (15,000 cells/well) and used at ≥80% confluence. Growth medium was replaced, and cells were pre-treated with vehicle or antagonists for 30 min, followed by stimulation with agonists or positive controls for 6 or 24 h. Supernatants (150 µL) were collected and stored at −80 °C until analysis.

Cytokines were measured using the Bio-Plex human screening panels according to the manufacturer’s instructions. Fluorescence intensities were converted to cytokine concentrations (pg/mL) based on standard curves generated using Bio-Plex Manager Software v6.2. Values below or above the detection limits were assigned the minimum or maximum quantifiable concentration, respectively.

## 3. RESULTS

### 3.1 H1R is the primary mediator of histamine-induced Ca²⁺ signaling in LECs

Histamine is an endogenous agonist that activates four G protein-coupled receptor (GPCR) subtypes: H1R – H4R [19, 20, 29], each with distinct G protein coupling and downstream signaling profiles [30]. The functional expression of histamine receptor subtypes in LECs was assessed based on intracellular Ca^2+^ responses to histamine in combination with subtype-selective antagonists.

LECs exhibited a concentration-dependent response to histamine, characterized by a rapid increase in intracellular Ca²⁺ followed by a sustained plateau phase **(Fig. 1A)**. To dissect the contribution of each receptor subtype, cells were pre-treated with the following selective antagonists: diphenhydramine (H1R) **(Fig. 1B)**, ranitidine (H2R) **(Fig. 1C)**, S38093-2 (H3R) **(Fig. 1D)**, and JNJ7777120 (H4R) **(Fig. 1E)**. The effects of each antagonist on histamine-evoked Ca^2+^ signaling are presented as kinetic traces **(Fig. 1B-E)** and concentration-response curves **(Fig. 1F)**.

**Figure 1.**
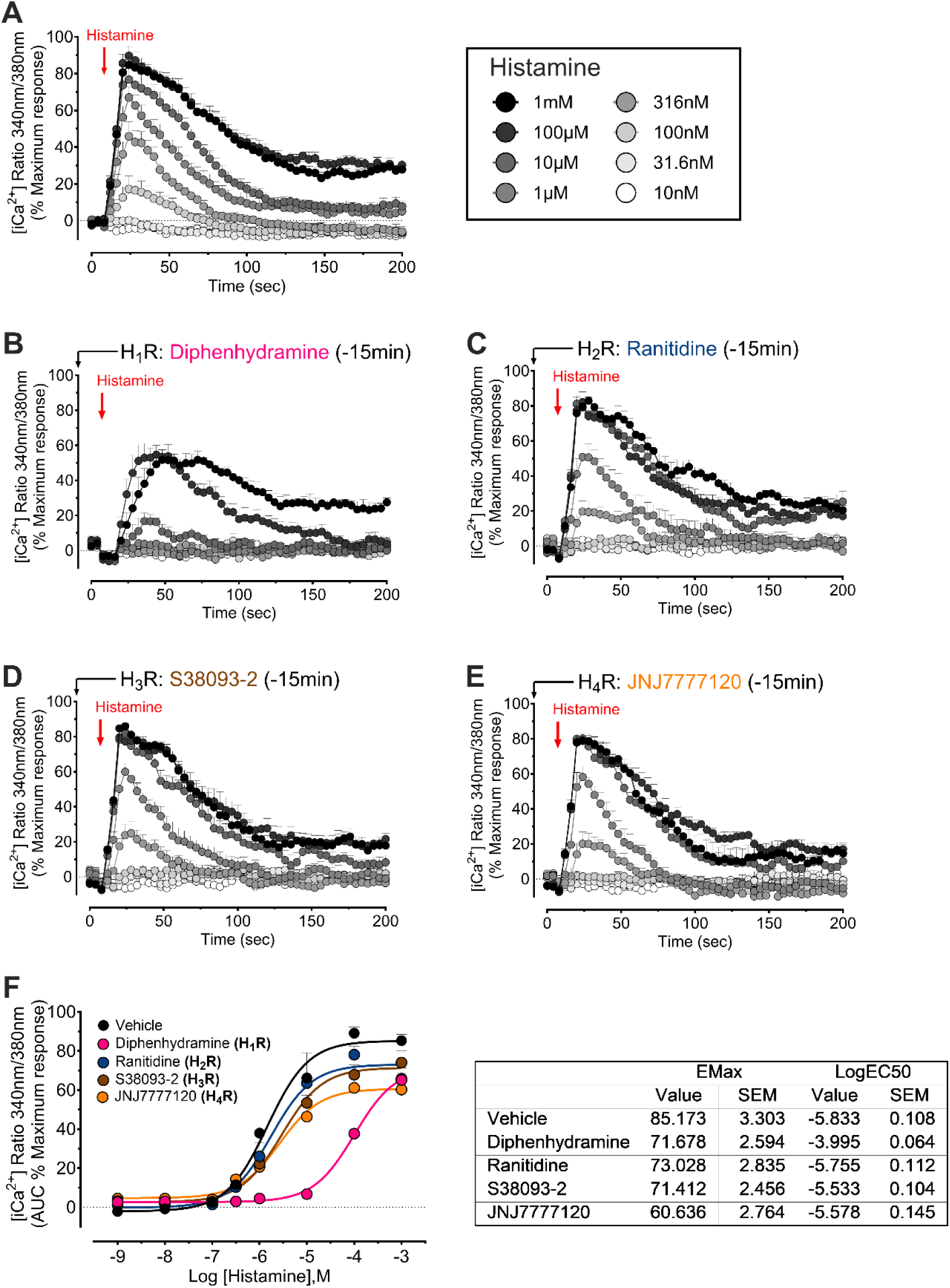
Histamine induced calcium mobilization in primary LECs. Ca^2+^ traces for LECs stimulated with increasing concentrations of histamine following pretreatment with vehicle (**A**) or antagonists selective for (**B**) H_1_R (Diphenhydramine, 1µM), (**C**) H_2_R (Ranitidine, 1µM), (**D**) H_3_R (S38093-2, 1µM) or (**E**) H_4_R (JNJ7777120, 1µM). (**F**) Concentration response curves summarizing effects of antagonists on histamine-evoked Ca^2+^ signaling (area under the curve for time traces shown in panel **A-E**). Data are presented as mean ± S.E.M., n=5 independent replicates.

H1R inhibition with diphenhydramine (1 µM) produced the most pronounced effect on histamine-evoked Ca²⁺ signaling, resulting in an approximately 30-fold reduction in potency (pEC_50_) and a measurable decrease in maximal response (Emax) **(Fig. 1)**. These findings are consistent with previous reports that H1R is functionally expressed by LECs [22] and with the known coupling of H1R to G_q_ proteins, which mediate phospholipase C activation and intracellular Ca²⁺ mobilization [20]. In comparison, blockade of H2R with ranitidine (1 µM) and H3R with S38093-2 (1 µM) only modestly reduced the Emax without affecting histamine potency **(Fig. 1F)**. In addition to diphenhydramine, the most pronounced effect on efficacy was observed with H4R blockade using JNJ7777120 (1 µM), which reduced the maximal Ca²⁺ response but did not alter potency **(Fig. 1F)**.

Together, these results indicate that H1R serves as the primary receptor subtype mediating histamine-evoked Ca²⁺ signaling in LECs. Importantly, the strong reduction of Emax following H4R inhibition suggests a secondary but significant contribution of H4R to the overall Ca²⁺ response, whereas H2R and H3R make only minor contributions.

### 3.2 TRPV4 partly mediates histamine-evoked Ca²⁺ responses in LECs

Activation of histamine receptors increases intracellular Ca²⁺ levels either through release from intracellular stores or by channel-dependent influx from the extracellular environment [20, 22, 31]. Previous studies have identified members of the TRP channel family, including TRPV4, as mediators of histamine-induced Ca²⁺ influx [25–27, 32]. We therefore examined the contribution of TRPV4 to histamine-evoked Ca²⁺ signaling in LECs.

First, we confirmed previous reports that LECs express functional TRPV4 [10, 33, 34] using the potent TRPV4 agonist GSK1016790A (GSK101). GSK101 produced a robust, concentration-dependent Ca²⁺ response (**Fig. 4A**). The contribution of extracellular Ca²⁺ influx to histamine receptor signaling was determined by removal of free extracellular Ca²⁺ with EGTA (2 mM). This resulted in significant reductions to both the potency and amplitude of histamine-evoked responses **(Fig. 2A, C, D)**. LECs were pretreated with the selective TRPV4 antagonist HC-067047 (HC067, 10 µM) to directly test the involvement of TRPV4. HC067 significantly attenuated histamine-induced Ca²⁺ responses, reducing both Emax and pEC_50_ **(Fig. 2B, C, D)**. The inhibitory effect of HC067 closely mirrored that of extracellular Ca²⁺ removal (HC067 pEC₅₀: 5.073; EGTA pEC₅₀: 5.096), suggesting that influx via TRPV4 contributes to elevated intracellular Ca²⁺ in response to histamine receptor signaling.

**Figure 2.**
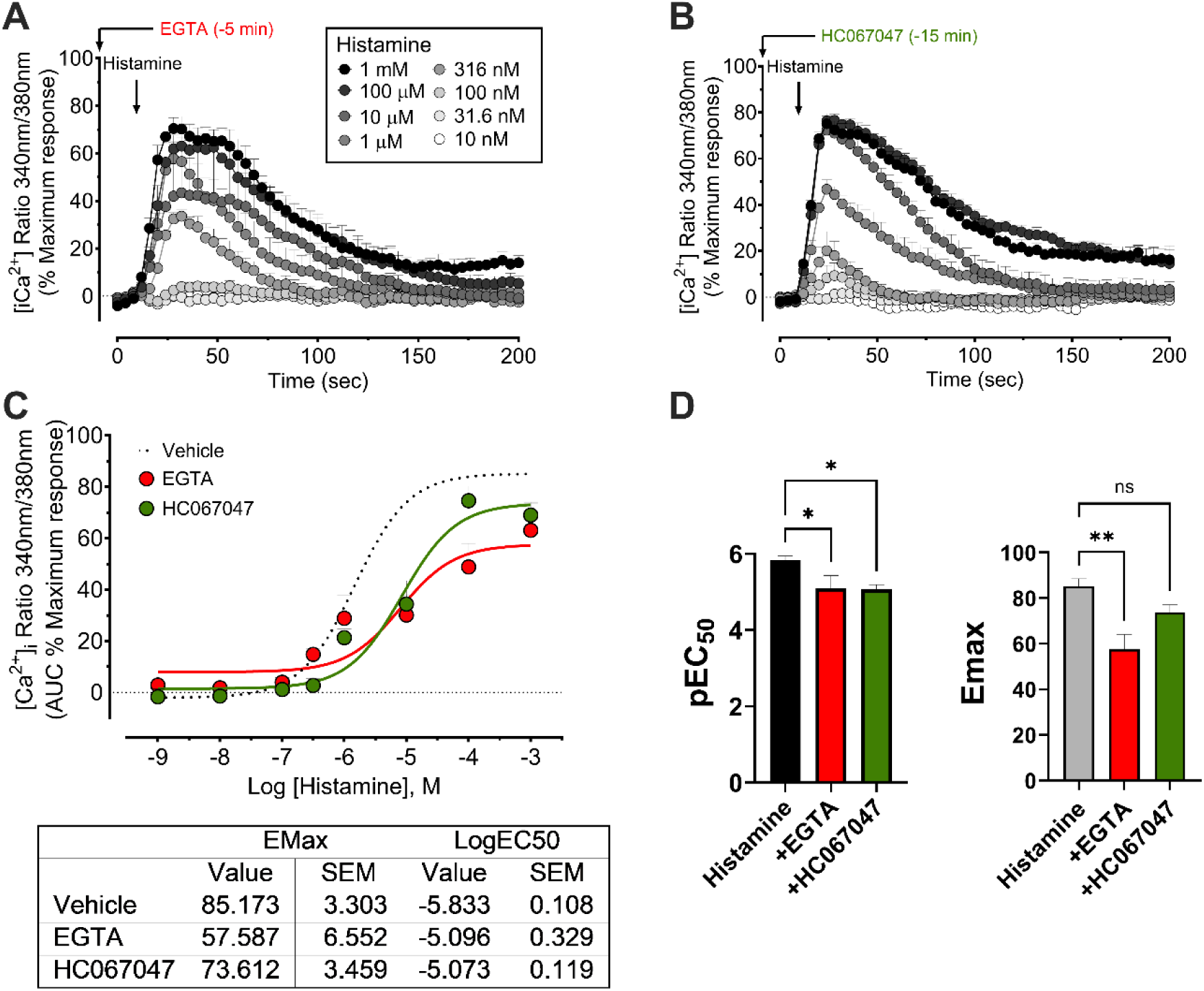
TRPV4 contributes to histamine-evoked calcium responses in LECs. **A.** Time traces showing Ca^2+^ responses by LECs to increasing concentrations of histamine. Responses were attenuated following pretreatment with a calcium chelator (EGTA, 2 mM) (**A**) or the TRPV4 antagonist HC067 (10 µM) (**B**). **C**. Concentration response curves based on area under the curve for traces shown in A and B. **D**. pEC50 and Emax values from concentration response curves shown in A and B. Mean ± S.E.M., n=5 independent experiments in duplicate.

TRPV4 promotes Ca²⁺ signaling downstream of the G_i/o_-coupled H4R subtype in primary keratinocytes [27]. Based on this study, we investigated H4R–TRPV4 crosstalk in LECs. The selective H4R agonist 4-methylhistamine (4-MeHA) [35] elicited a concentration-dependent Ca²⁺ response that was smaller in amplitude relative to Ca^2+^ signaling to histamine (pEC_50_: 5.833 vs 5.062; Emax: 49.46 vs 85.17) **(Fig. 3A)**. Responses to 4-MeHA were markedly reduced by either extracellular Ca²⁺ removal (EGTA, 2 mM) **(Fig. 3B)** or TRPV4 blockade (HC067) **(Fig. 3C)** and were abolished by the H4R-selective antagonist JNJ1777120 (1 µM), confirming H4R dependence **(Fig. 3D)**.

**Figure 3.**
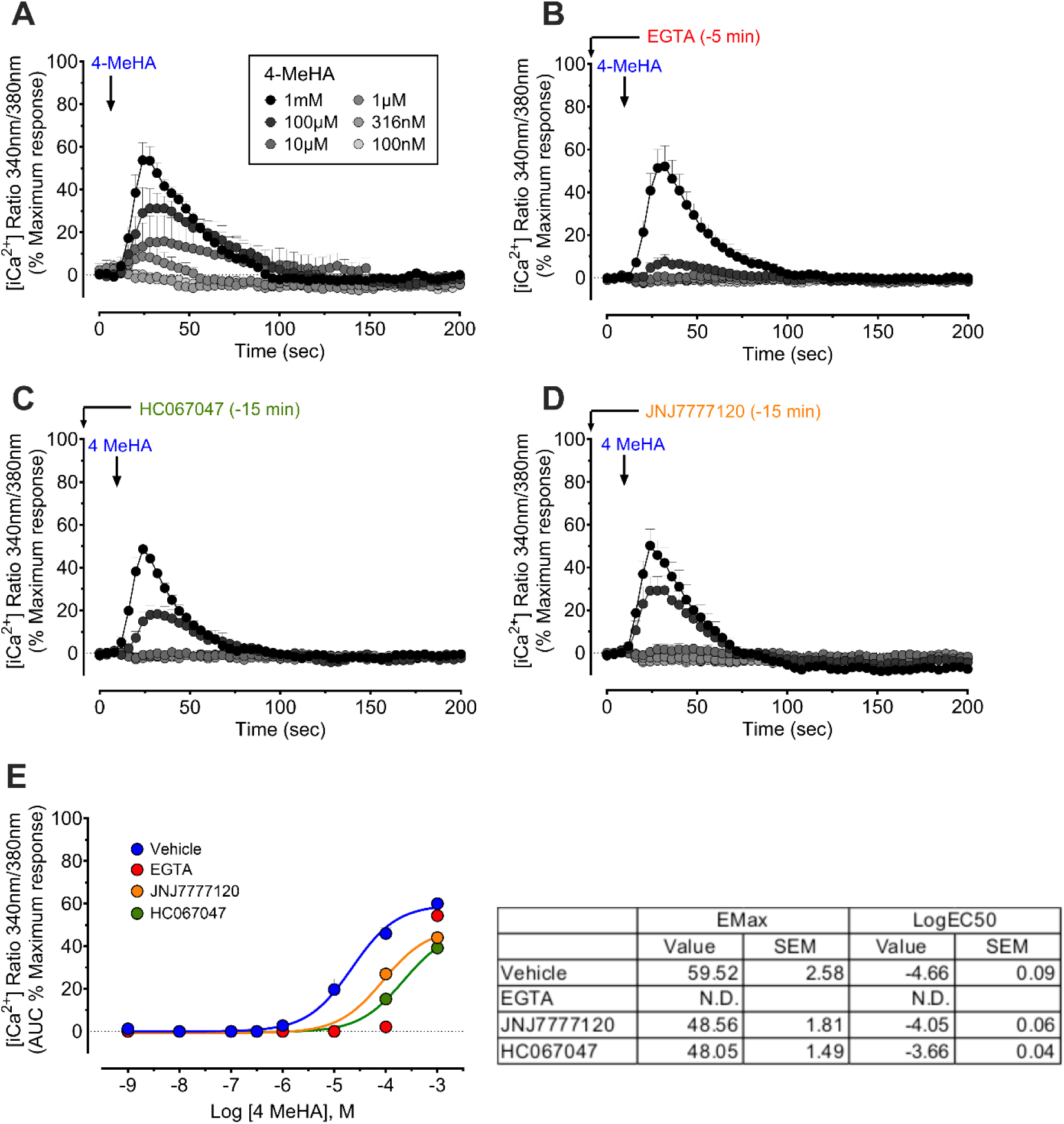
H_4_R-evoked calcium responses in LECs are TRPV4-dependent. **A.** Ca^2+^ traces showing concentration-dependent responses to the H_4_R selective agonist 4-MeHA. The response to 4-MeHA was attenuated following pretreatment of LECs with the Ca^2+^ chelator EGTA (2mM) (**B**), the TRPV4 antagonist HC067 (10µM) (**C**), or the H_4_R antagonist JNJ7777120 (1µM) (**D**). **E**. Concentration response curves summarising data presented in panels A-D (area under the curve). Data are presented as mean ± S.E.M., n=3-5 independent experiments in duplicate.

Collectively, these data support a model in which TRPV4 contributes significantly to histamine-evoked Ca²⁺ influx in LECs downstream of both H_1_R and H4R subtypes.

### 3.3 TRPV4 signaling in LECs is enhanced by histamine receptor activation via a PLA2-dependent mechanism

TRPV4 activity can be sensitized by prior activation of GPCRs, leading to enhanced responsiveness to physical and chemical stimuli [13]. Previous studies have demonstrated that histamine receptor activation augments subsequent TRP channel signaling, implicating this mechanism in pathophysiological processes such as pain and inflammation [25, 26, 32, 36].

The effect of histamine receptor activation on TRPV4 signaling in LECs was assessed. Histamine pretreatment (1 µM, 15 min) caused a modest leftward shift in the GSK101 concentration–response curve (pEC₅₀: vehicle = 6.51; histamine = 7.81) along with a reduction in Emax relative to vehicle pretreated cells **(Fig. 4A, B, D)**. Notably, pretreatment with 4-MeHA (1 µM, 15 min) resulted in greater TRPV4 sensitization, corresponding to an 89-fold increase in potency (pEC₅₀ from 6.51 to 8.46) **(Fig. 4C, D)**. These findings suggest that H4R activation enhances TRPV4 sensitivity to agonists, supporting functional crosstalk between these pathways in LECs.

**Figure 4.**
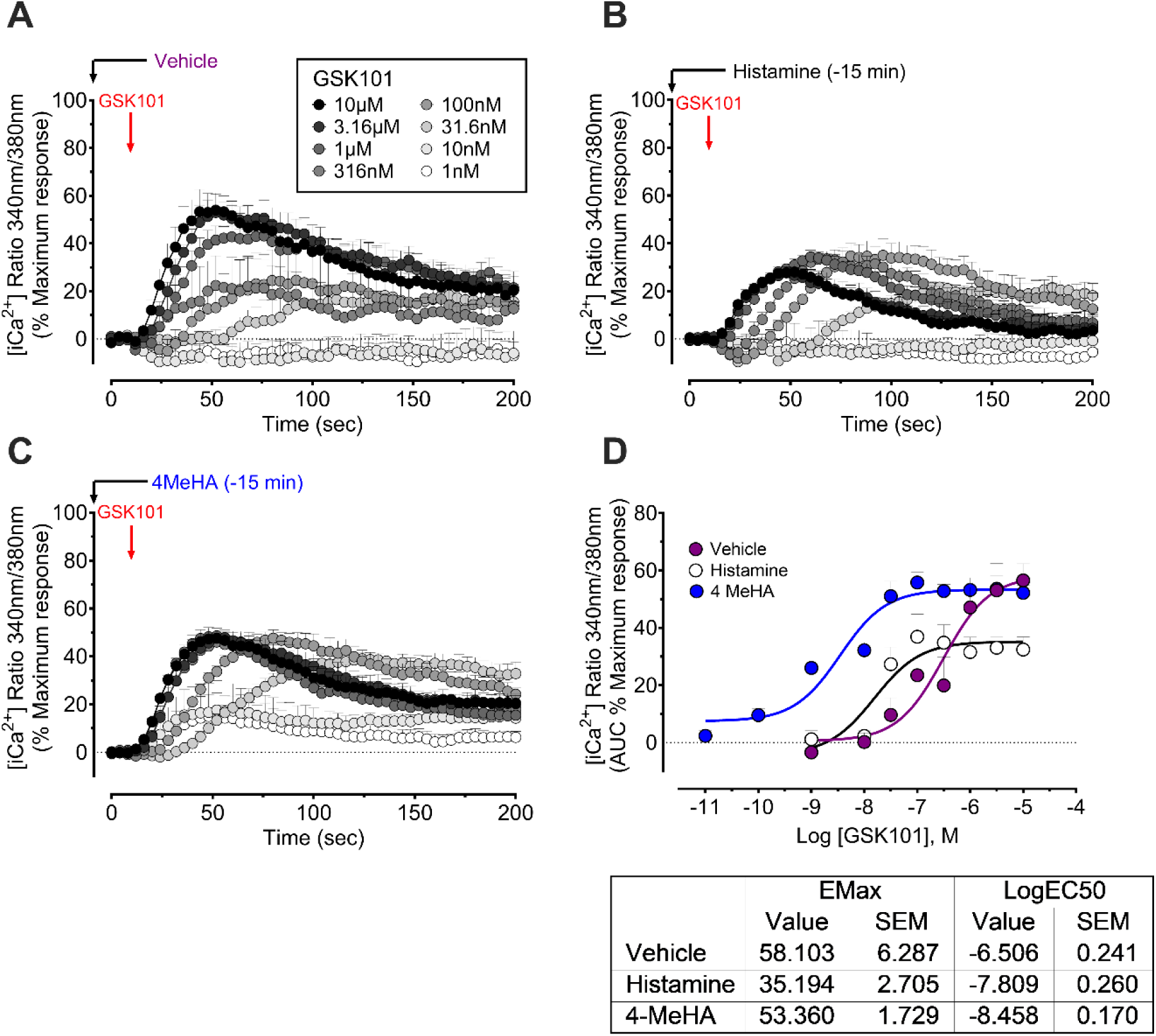
Histamine receptor mediated sensitization of TRPV4 in LECs. Ca^2+^ traces for LECs stimulated with increasing concentrations of the TRPV4 selective agonist GSK101 (**A**) and equivalent responses following pretreatment with histamine (1µM, **B**) or 4-MeHA (1µM, **C**). (**D**) Concentration response curves showing the effects of histamine receptor stimulation on Ca^2+^ responses to GSK101 (area under the curve for time traces shown in panel **A-C**). (**E**) Table summarising the pEC50 and Emax data presented in panel (**D**). Data are presented as mean ± S.E.M., n=3-5 independent experiments in duplicate.

Sensitization of TRPV4 downstream of GPCRs, including in response to histamine receptor activation [26], has been linked to phospholipase A2 (PLA2) activity [13, 15]. PLA2 catalyzes the generation of arachidonic acid from phospholipids, leading to production of endogenous TRPV4 agonists such as 5,6-EET [16]. To evaluate whether PLA2 mediates H4R-dependent TRPV4 potentiation in LECs, we tested inhibitors targeting cytosolic PLA2 (cPLA2) and secretory PLA2 (sPLA2) isoforms. Both inhibitors significantly reduced Ca²⁺ responses to 4-MeHA **(Fig. 5A)**, confirming that PLA2 activity is required for H4R Ca²⁺ signaling.

**Figure 5.**
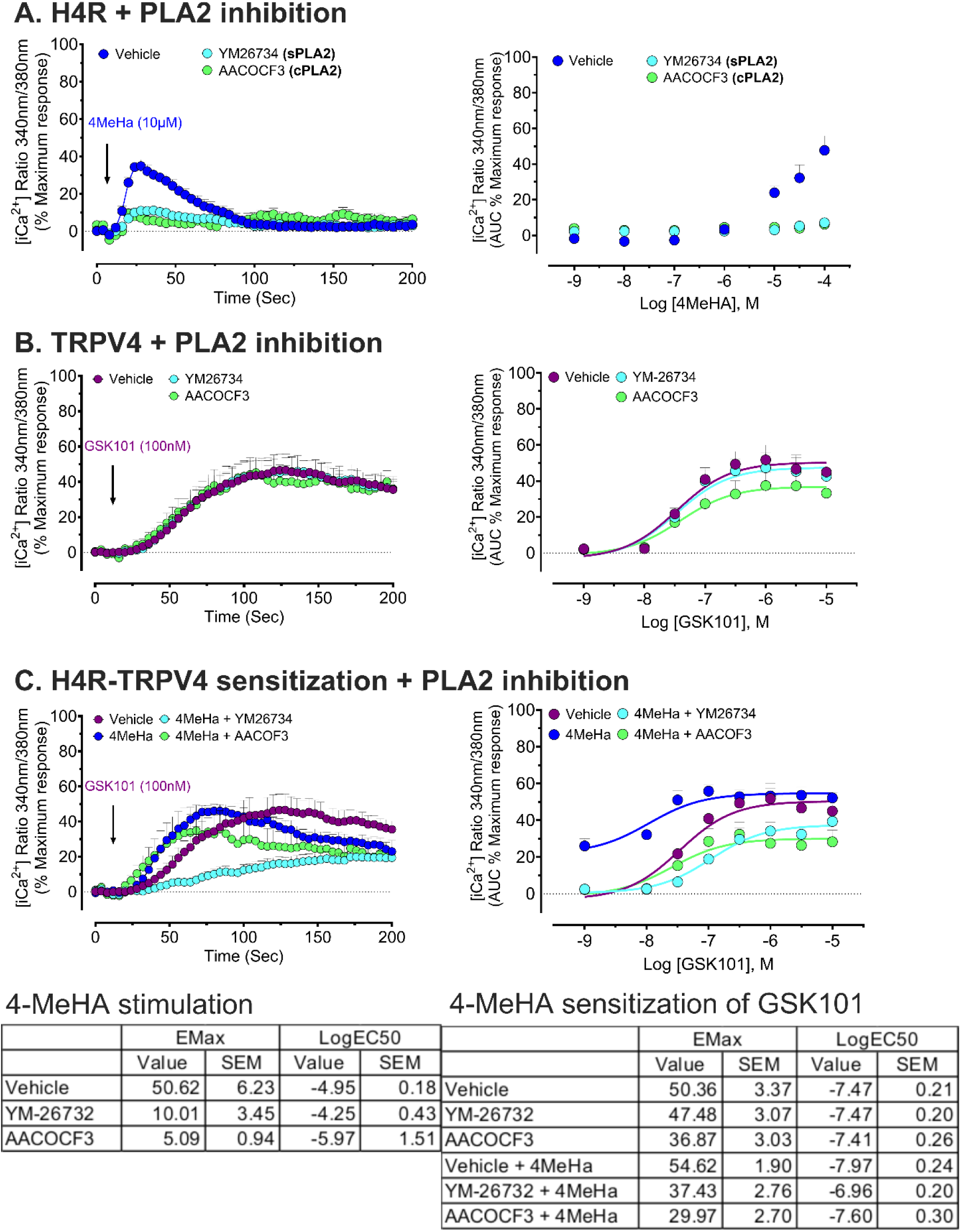
Ca^2+^ signaling in response to H_4_R activation and H_4_R-dependent sensitization of TRPV4 are mediated through PLA2 in LECs. **A.** Inhibition of PLA2 (YM-26734, 1µM or AACOCF3, 1µM) effectively blocked Ca^2+^ signaling to 4-MeHA, as shown by time traces (left) and concentration response curves (right). **B.** In contrast, PLA2 inhibition had minimal effect on TRPV4-dependent Ca^2+^ responses. **C.** Pretreatment of LECs with 4-MeHA resulted in enhanced responses to GSK101, which was mediated through a PLA2-dependent mechanism. Data are presented as mean ± S.E.M., n=3-6.

To determine whether basal PLA2 activity directly modulates TRPV4 function, we assessed responses to GSK101 in the presence of these inhibitors. The cPLA2 inhibitor AACOCF₃ caused a modest reduction in GSK101-evoked Ca²⁺ responses, whereas sPLA2 inhibition had no effect **(Fig. 5B)**. This observation indicates that while PLA2 may have a minor direct effect on TRPV4, its primary role is in receptor-operated channel activity. Importantly, both cPLA2 and sPLA2 inhibitors abolished 4-MeHA-mediated sensitization of TRPV4 **(Fig. 5C)**, confirming that PLA2 activity is essential for histamine receptor-dependent enhancement of TRPV4 activity.

Together, these results demonstrate that histamine receptors and TRPV4 are co-expressed and functionally coupled in LECs. This interaction is mediated through a PLA2-dependent mechanism, highlighting a signaling axis that amplifies Ca²⁺ responses downstream of inflammatory cues.

### 3.4 Histamine promotes NFAT translocation via TRPV4 in LECs

The nuclear factor of activated T cells (NFAT) is a calcium-regulated transcription factor that plays a critical role in lymphatic development and lymphangiogenesis [37]. Both GPCRs and TRPV4 have been implicated in the activation of the calcineurin–NFAT pathway, leading to transcriptional changes in endothelial and epithelial systems [31, 38]. To assess whether TRPV4 and histamine signaling promote NFAT activation in LECs, we evaluated the subcellular localization of NFAT following stimulation. LECs were treated with GSK101 or histamine receptor agonists and subsequently fixed and immunolabelled with an anti-NFATc1-2 specific antibody **(Fig. 6)**.

**Figure 6.**
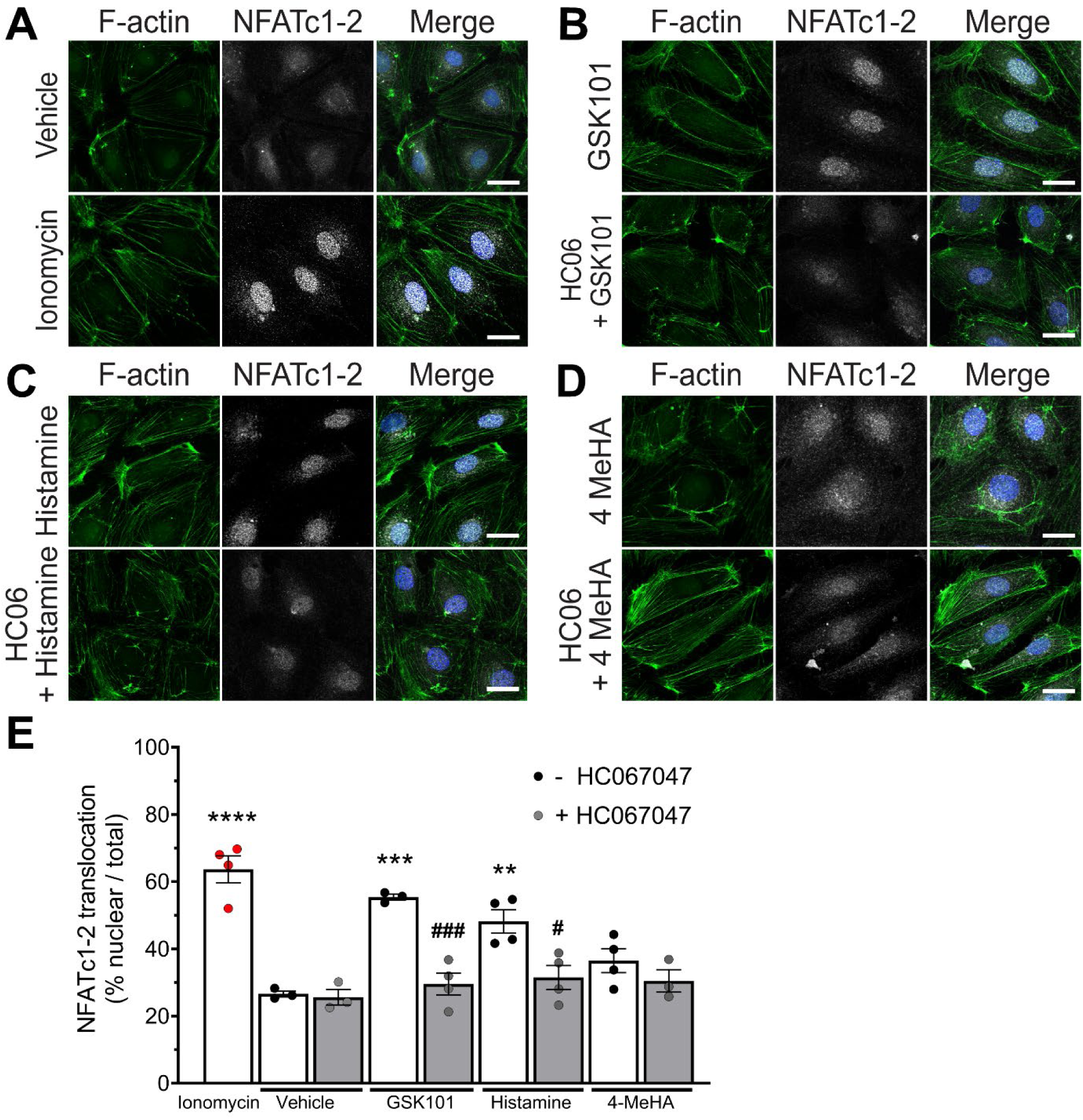
TRPV4 affects histamine-induced nuclear translocation of transcription factor NFATc1-2 in LECs. Immunofluorescence imaging of LECs stained with DAPI (nucleus, blue), F-actin (green) and the transcription factor NFATc1-2 (white) (**A-D**). LECs treated with the positive control ionomycin (1 µM, 30 min) have a higher nuclear/total NFATc1-2 ratio compared with vehicle (**A, E**). NFAT translocation imaging in LECs treated with GSK101 (100nM) (**B**), histamine (1 µM) (**C**) and 4-MeHA (1 µM) (**D**) after pre-treatment with vehicle or HC067. (**E**) Quantitative image analysis of NFAT translocation (% nuclear NFAT vs. total NFAT) for individual cells. Mean ± S.E.M., n= 50 cells, N=3-4 independent replicates. Statistical analysis: *** p≤ 0.001 compared to vehicle, ** p≤ 0.01 compared to vehicle, ### p≤0.001 compared to without HC067, # p≤0.05 compared to without HC067, one-way ANOVA Tukey’s multiple comparison.

Treatment with the calcium ionophore ionomycin (1 µM, 30 min) served as a positive control and induced robust translocation of NFAT from the cytoplasm into the nucleus compared to vehicle-treated cells **(Fig. 6A)**. Similarly, activation of TRPV4 with GSK101 (100 nM) promoted marked nuclear localization of NFAT. NFAT translocation was prevented by pre-treatment with HC067, confirming TRPV4 dependence **(Fig. 6B)**.

Treatment of LECs with histamine or 4-MeHA (1 µM, 30 min) led to a significant increase in nuclear NFAT transport **(Fig. 6C, D)**. Histamine receptor-evoked NFAT translocation to the nucleus was inhibited by HC067 pretreatment **(Fig. 6E)**, consistent with the requirement for TRPV4-mediated Ca^2+^ mobilization for this process.

These results demonstrate that histamine receptor activation leads to the nuclear translocation of NFAT via a TRPV4-dependent mechanism. While direct transcriptional activity was not assessed in this assay, the observed redistribution of NFAT strongly supports the role of TRPV4 as a downstream effector linking histamine signaling to Ca²⁺-dependent gene transcription and proliferation in LECs.

### 3.5 TRPV4 activation alters junctional architecture and cytoskeletal organization in LECs

LEC function relies on dynamic regulation of cell–cell junctions and the cytoskeleton to maintain vascular integrity and respond to inflammatory signals. To assess whether TRPV4 contributes to junctional remodeling and actin organization, we performed immunofluorescence imaging of the endothelial adherens junction protein VE-cadherin, F-actin, and the nuclear marker DAPI in LECs treated with GSK101 (100 nM) or histamine (1 µM), in the presence or absence of HC067 (10 µM).

TRPV4 activation with GSK101 resulted in prominent remodeling of VE-cadherin at cell–cell borders from a discontinuous and jagged appearance under basal conditions to continuous linear staining (**Fig. 7A, B**, left panels). In parallel, GSK101 induced a marked reorganization of F-actin into prominent cortical bundles and stress fibers (**Fig. 7B**, middle panel), indicative of cytoskeletal tension. These changes were accompanied by a loss of well-defined junctions and altered cellular morphology in merged images (**Fig. 7A, B**, right panels). Pre-treatment with HC067 prevented both VE-cadherin disassembly and actin remodeling, confirming TRPV4-dependence of these structural changes (**Fig. 7C**). In contrast, histamine treatment led to increased formation of stress fibers without apparent disruption of VE-cadherin localization (**Fig. 7D**). F-actin accumulated centrally, forming dense transverse bundles across the cytoplasm, while VE-cadherin remained largely continuous at cell borders. These changes were not prevented by HC067 (**Fig. 7E**), suggesting that histamine-induced actin remodeling occurs through a TRPV4-independent pathway. Notably, histamine did not replicate the junctional remodeling phenotype observed with direct TRPV4 activation, reinforcing the selective role of TRPV4 in modulating endothelial barrier integrity.

**Figure 7.**
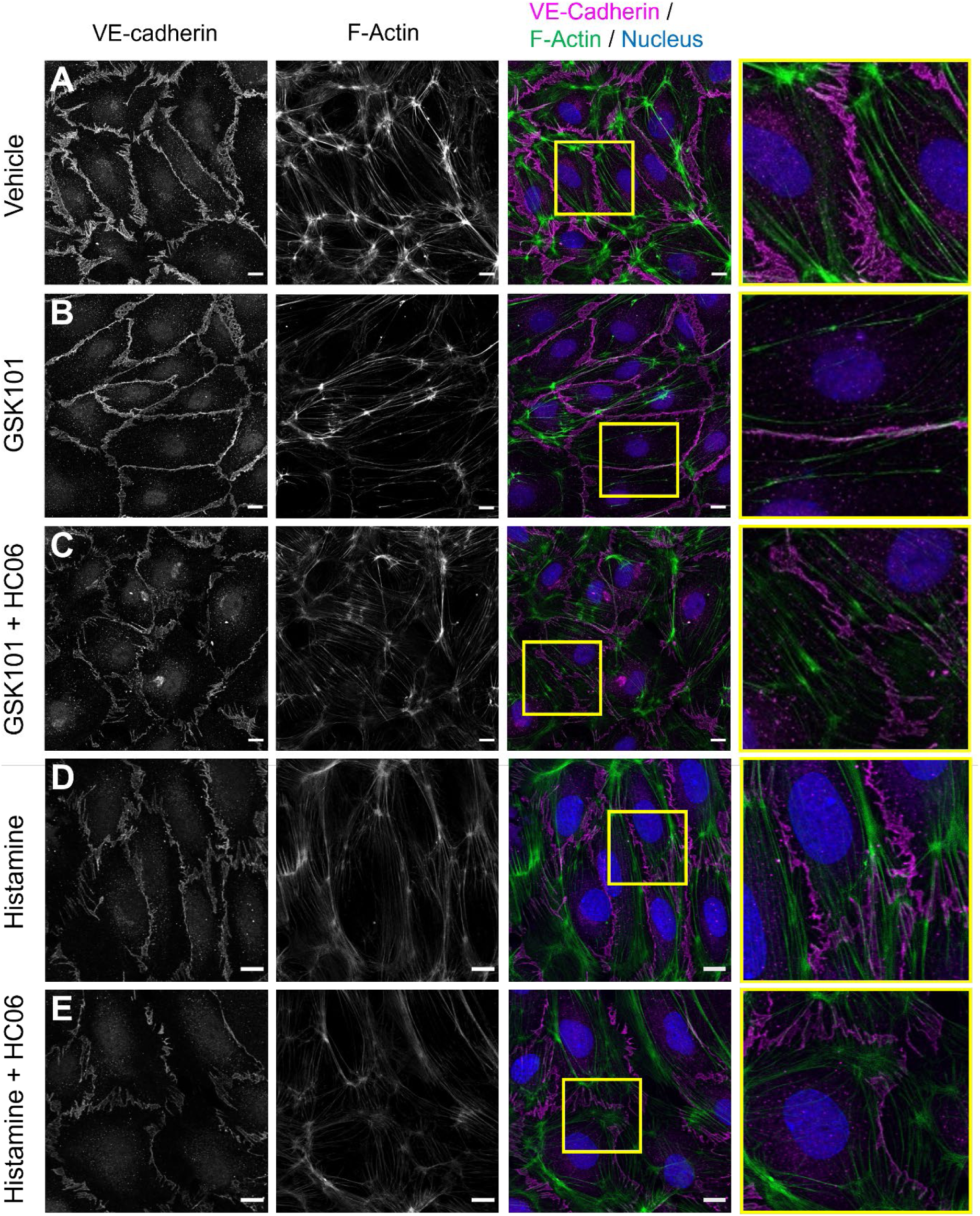
TRPV4- and histamine receptor-induced junctional and cytoskeletal remodeling in LECs. Immunofluorescence imaging of LECs stained for the nuclear marker DAPI (blue), F-actin (green) and VE-cadherin (magenta). LECs were treated with: (**A**) vehicle, (**B**, **C**) GSK101 (100nM, 30 min) with or without the TRPV4 inhibitor HC067 (10 µM), or (**D**, **E**) histamine (1 µM, 30 min) with or without HC067. Activation of TRPV4 led to extensive changes to LEC junctions and to cytoskeletal remodeling. Representative images from n=3 independent experiments performed in duplicate. Scale: 10 µm.

Together, these results demonstrate that TRPV4 activation leads to coordinated junctional and cytoskeletal remodeling in LECs, whereas histamine induces more selective changes to the actin cytoskeleton. These observations support a model in which TRPV4 acts as a transducer of mechanical and chemical signaling, leading to regulation of LEC structural dynamics.

### 3.6 Histamine receptors and TRPV4 promote selective cytokine release from LECs

Histamine promotes immune cell chemotaxis not only by inducing physiological changes such as smooth muscle contraction and vasodilation, but also by stimulating cytokine production by immune and endothelial cells [20, 22, 31]. LECs are increasingly recognized as active participants in immune regulation [39, 40] and can produce a range of cytokines and growth factors [41]. Given that TRPV4-mediated Ca²⁺ influx promotes nuclear translocation of NFAT in LECs, we next investigated whether histamine receptor and TRPV4 activation could stimulate downstream secretion of inflammatory mediators.

The effects of histamine and TRPV4 activation on cytokine production by LECs were quantified using a Bio-Plex™ assay. Cells were treated with histamine (10 µM), GSK101 (100 nM), or 4-MeHA (10 µM), and supernatants were collected after 6 and 24 h. PDBu (100 nM) and ionomycin (1 µM) were used as positive controls.

TRPV4 activation with GSK101 robustly increased secretion of CCL4 (MIP-1β) and CCL5 (RANTES) at both 6 and 24 h, as well as IL-9 at 24 h. This finding supports a TRPV4-specific role in regulating the release of these chemokines **(Fig. 8)**. Treatment with either histamine or 4-MeHA did not induce secretion of any cytokines tested, suggesting that H1R and H4R activation alone is insufficient to trigger measurable cytokine release under these conditions **(Fig. 8)**. Whether histamine or 4-MeHA pretreatment could enhance TRPV4-evoked chemokine release was not assessed.

**Figure 8.**
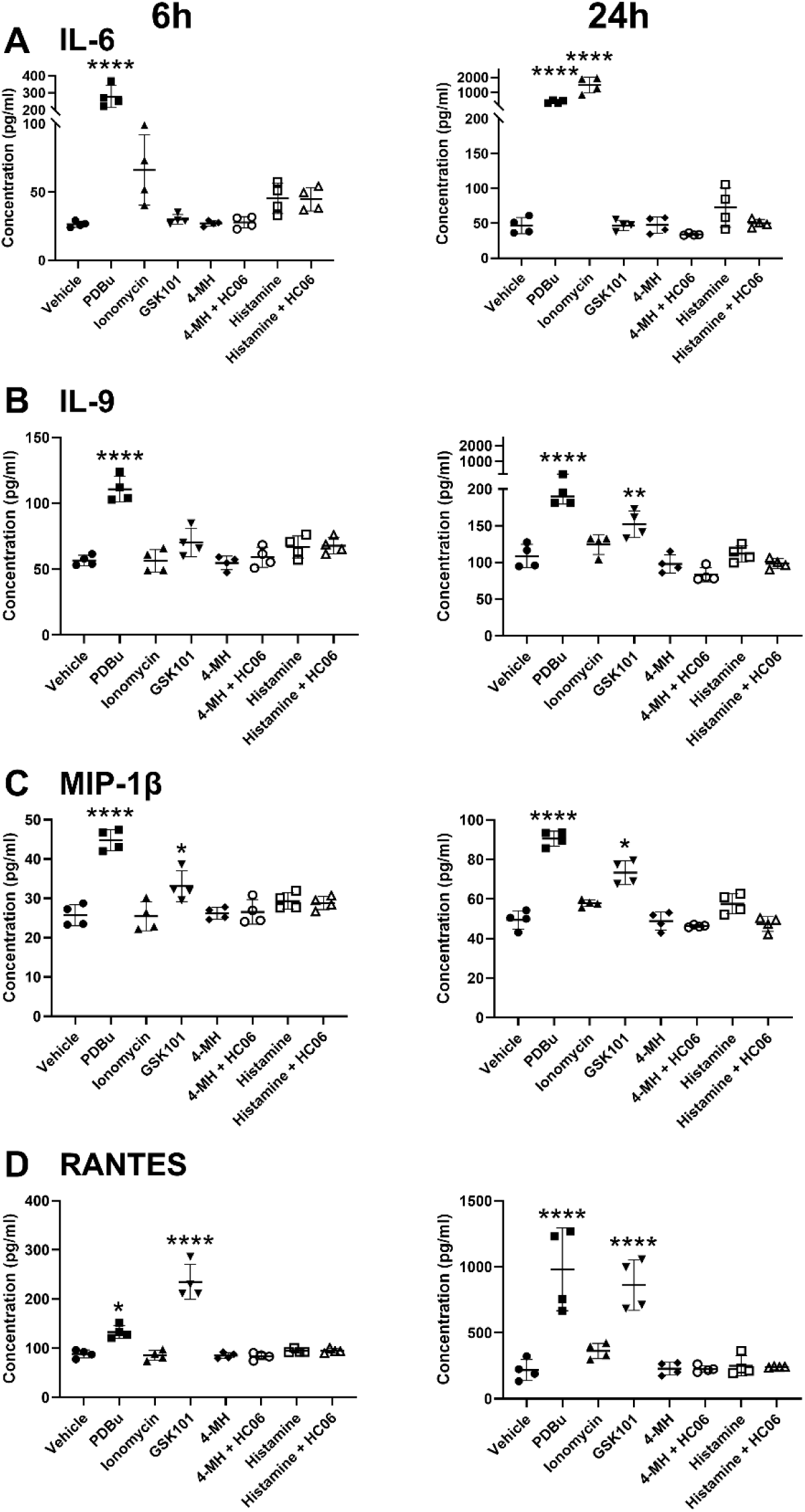
TRPV4 activation increased secretion of inflammatory cytokines IL-9, CCL4, and CCL5 in LECs. GSK101 induced secretion of cytokines MIP-1β (CCL4) and RANTES (CCL5) in LECs at both 6 and 24 h time points and IL-9 at 24 h. Treatment of LEC s with histamine or 4-MeHA did not have a significant effect on the secretion of the selected cytokines at either time point. Data are presented as mean ± S.E.M., n=4 independent replicates. Statistical analysis: * p < 0.05, ** p <0.01, ****; p < 0.0001 compared to vehicle, one-way ANOVA with Tukey’s multiple comparison test.

No significant changes were observed in the secretion of other interleukins (IL-4, IL-5, IL-10, IL-15, IL-17) or growth factors (GM-CSF, G-CSF, PDGF, VEGF) at either time point **(Supplementary Figs S1 and S2)**. Additionally, classical pro-inflammatory cytokines such as IL-1β, IL-2, TNF-α, and IFN-γ were unaffected by histamine, GSK101, or 4-MeHA treatments **(Supplementary Fig. S3)**.

Together, these findings demonstrate that activation of TRPV4 can selectively promote cytokine release in LECs. Notably, the two most prominent cytokines that were upregulated by TRPV4 activation (CCL4 and CCL5) are chemotactic proteins involved in immune cell migration [42, 43].

## 4. DISCUSSION

The lymphatic endothelium plays a critical role in maintaining tissue fluid homeostasis, modulating immune surveillance, and responding to inflammatory cues. While Ca²⁺ signaling is recognized as a key regulator of LEC function, the molecular mechanisms that couple extracellular stimuli to Ca²⁺-dependent processes in LECs remain incompletely understood. In this study, we identify a novel H4R-PLA2-TRPV4-mediated Ca²⁺ signaling axis in primary human dermal LECs that regulates not only Ca²⁺ dynamics, but also downstream transcriptional, morphological, and secretory responses.

Histamine exerts pleiotropic effects via four distinct GPCR subtypes (H1R-H4R), each with unique G protein coupling and signaling profiles [19, 20]. While H1R is classically linked to G_q_-IP₃-Ca²⁺ release from stores, H4R is G_i/o_-coupled and has been associated with chemotaxis, actin remodeling, and immune cell function [44, 45]. We confirm that both H1R and H4R are functionally expressed by LECs, with H1R primarily responsible for mobilizing Ca²⁺. We demonstrate that H4R can also promote elevations in intracellular Ca²⁺ levels via a non-canonical mechanism involving TRPV4 activation.

Our data reveal that H4R activation by 4-MeHA enhances TRPV4 sensitivity to the agonist GSK101, resulting in an ∼89-fold increase in potency. This sensitization was abolished by inhibition of cytosolic and secretory PLA2 isoforms, consistent with H4R-dependent activation of PLA2 and subsequent production of endogenous TRPV4 agonists such as epoxyeicosatrienoic acids (EETs) [15, 26]. These findings position TRPV4 as a downstream effector of a G_i/o_-coupled GPCR, which differs from the more commonly associated involvement of TRP channels in G_αq_-mediated signaling [13].

Functionally, activation of this H4R-TRPV4 signaling pathway in LECs was associated with a diverse range of cellular responses. TRPV4 activation induced junctional remodeling and reorganization of the actin cytoskeleton, consistent with the established role of this ion channel in modulating endothelial barrier function. These effects were distinct from those induced by histamine, which promoted stress fiber formation without major disruption to the apparent integrity of VE-cadherin positive junctions, suggesting that TRPV4 activation triggers unique morphological changes. Additionally, TRPV4-mediated Ca²⁺ entry was required for nuclear translocation of NFATc1, a transcription factor implicated in lymphangiogenesis and LEC proliferation [37]. Importantly, NFATc1-translocation in response to histamine was prevented by TRPV4 inhibition, highlighting the role of TRPV4 in coupling GPCR signals to gene transcription. These observations differ to those of Si *et al.* 2020 [22], who reported that treatment of LECs with histamine led to disruption of VE-cadherin junctions through a calcium release activated channel (CRAC)-dependent mechanism, as confirmed by immunofluorescent labeling, increased macromolecule permeability, and reduced transendothelial electrical resistance. It should be noted that we did not assess monolayer permeability and therefore cannot exclude any effects of histamine or TRPV4 on junctional integrity. The requirement for Ca^2+^ entry via CRAC for histamine-evoked NFAT translocation has been demonstrated in HUVEC [31]. The similarities and differences between these studies and our own suggest that there is some redundancy in the coupling of histamine receptors to Ca^2+^ permeable channels, with both TRPV4 and CRAC capable of promoting equivalent downstream signaling.

Additional evidence for the functional divergence of histamine-evoked and TRPV4-mediated signaling was provided by the associated effects on cytokine release. Under our experimental conditions, activation of TRPV4 stimulated secretion of IL-9, CCL4 (MIP 1β), and CCL5 (RANTES), which are chemokines associated with T cell recruitment, transendothelial migration, and lymphangiogenesis [41, 46]. The observation contrasts with the lack of effect of histamine or 4-MeHA on cytokine release, suggesting that crosstalk between histamine receptors and TRPV4 is not sufficient to drive this process under basal conditions. Whether this interaction is changed under pathological conditions, such as inflammation, remains to be determined. Collectively, the findings of the present study suggest that TRPV4 may play a dual role in inflammatory signaling by LECs, with involvement in mediating both endothelial remodeling and immune cell recruitment, possibly through paracrine and autocrine signaling loops. The expression of CCR5, the receptor for CCL4 and CCL5, by both LECs and immune cells [47] further supports this hypothesis.

Together, our study provides compelling evidence that histamine receptors, particularly H4R, can transactivate TRPV4 via a PLA2-dependent mechanism in human LECs, resulting in significant alterations to cell signaling, shape, and secretory phenotype. This positions TRPV4 as a key effector node downstream of diverse GPCRs and a potential regulator of lymphatic endothelial responses to inflammation.

---

**Supplementary Figure 1.**
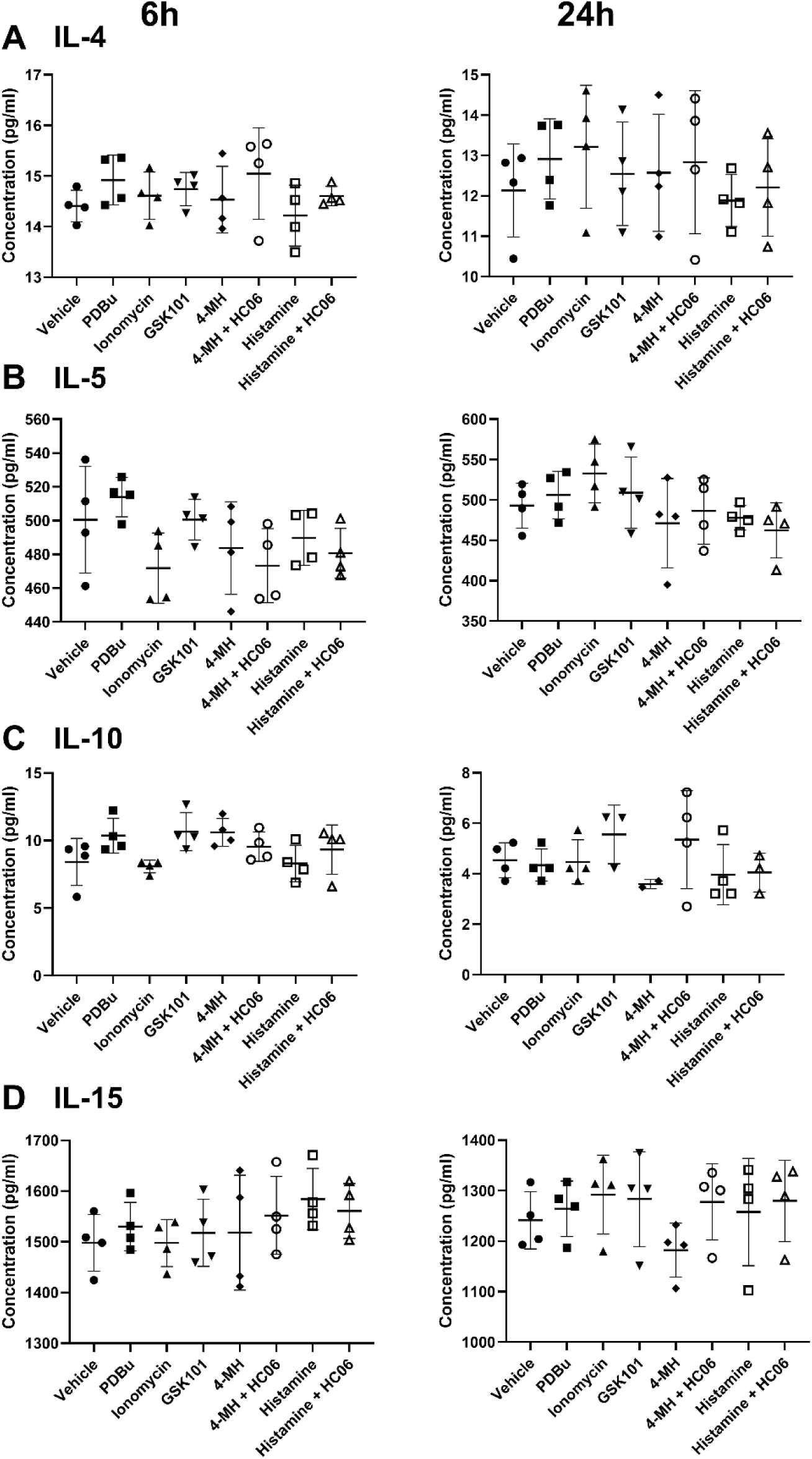
Secretion of interleukins IL-4, IL-5, 1L-10, and IL-15 is not affected by activation of TRPV4 or histamine receptors in LECs. Treatment of LECs with GSK101, histamine, or 4-MeHA did not promote secretion of these interleukins at either 6h or 24h. Data were analyzed by one-way ANOVA with Tukey’s multiple comparison test. Data are presented as mean ± S.E.M., n=4 independent replicates.

**Supplementary Figure 2.**
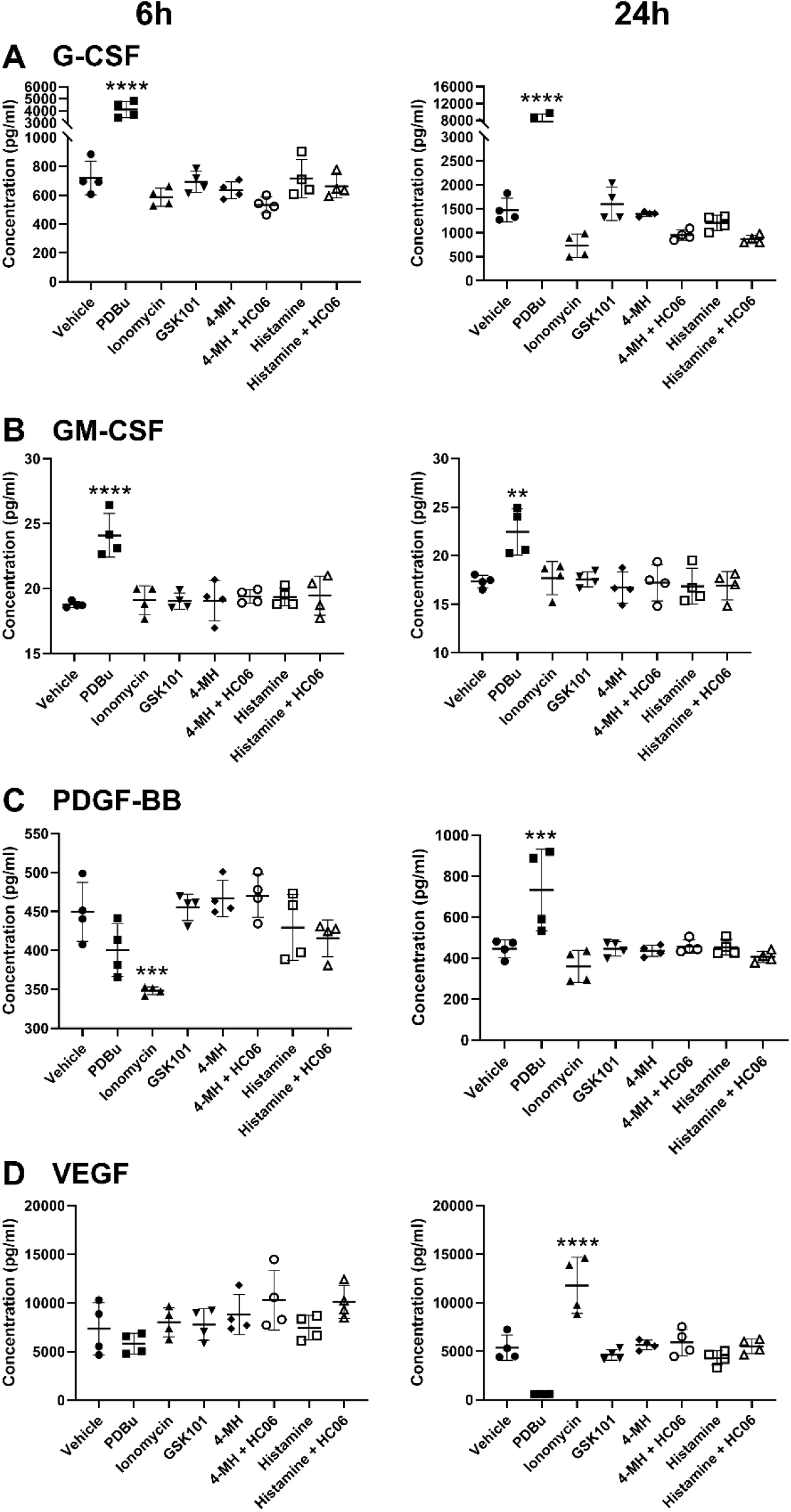
Secretion of the factors G-CSF, GM-CSF, PDGF-BB, and VEGF is not affected by activation of TRPV4 or histamine receptors in LECs. Treatment of LECs with GSK101, histamine, or 4-MeHA did not promote secretion of these factors at either 6h or 24h. Data were analyzed by one-way ANOVA with Tukey’s multiple comparison test. Data are presented as mean ± S.E.M., n=4. Statistical analysis: ** p <0.01, *** p <0.001, ****; p < 0.0001 compared to vehicle, one-way ANOVA with Tukey’s multiple comparison test.

**Supplementary Figure 3.**
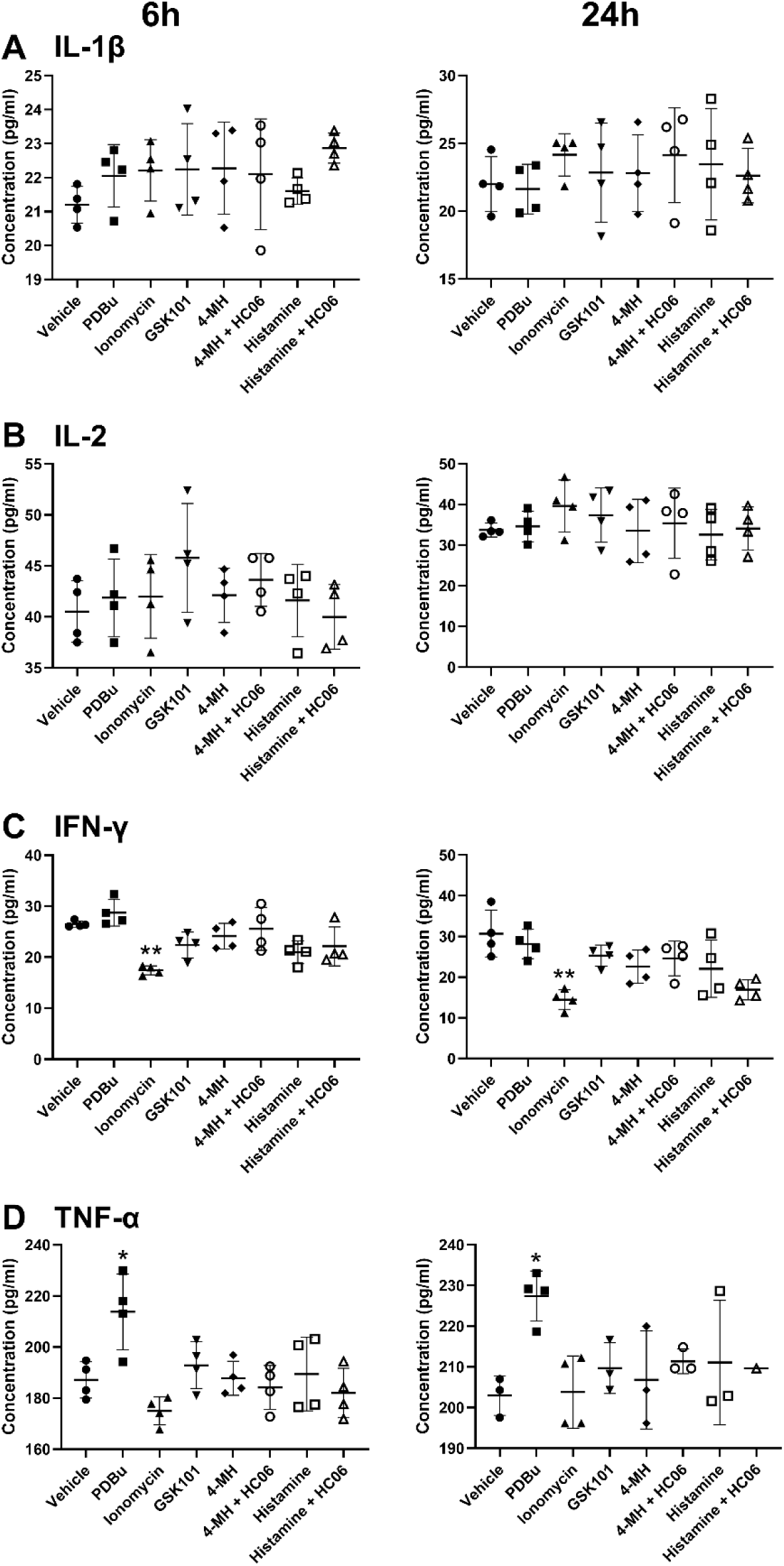
Secretion of the inflammatory mediators IL-1β, IL-2, IFN-γ, and TNF-α was not affected by activation of TRPV4 or histamine receptors in LECs. Treatment of LECs with GSK101, histamine, or 4-MeHA did not promote secretion of these mediators at either 6h or 24h. Data were analyzed by one-way ANOVA with Tukey’s multiple comparison test. Data are presented as mean ± S.E.M., n=4. Statistical analysis: * p<0.05, ** p <0.01 compared to vehicle, one-way ANOVA with Tukey’s multiple comparison test.

